# Phylogeny, ecology, and allometry: hierarchical controls of insect elemental composition

**DOI:** 10.64898/2025.12.11.693773

**Authors:** Léo Laborieux, Richard J. Knecht, William F. Fagan, Nathan J. Sanders, Anshuman Swain

**Affiliations:** Department of Ecology and Evolutionary Biology, University of Michigan, Ann Arbor, MI, USA; Museum of Paleontology, University of Michigan, Ann Arbor, MI, USA; Department of Biology, University of Maryland, College Park, MD, USA

**Keywords:** stoichiometry, evolutionary ecology, phosphorus, nitrogen

## Abstract

Elemental composition links organismal physiology to biogeochemical cycles, yet the relative importance of evolutionary and ecological drivers remains unresolved for insects. Integrating a global stoichiometric database with a comprehensive phylogenetic backbone, we demonstrate hierarchical control of insect stoichiometry, with phylogeny accounting for ∼50% of variation in C:N:P ratios, followed by temperature, body mass, trophic group and habitat. We also identify nutrient enrichment in terrestrial lineages relative to aquatic ones. Our work offers a general framework for partitioning evolutionary, ecological, and physiological influences on organismal stoichiometry, highlighting new avenues to link macroevolutionary patterns to underlying processes.

## Main

The elemental composition of life provides a universal currency for linking organismal biology to ecosystem processes, yet its ultimate determinants remain insufficiently understood (Sterner and Elser, 2002; Sardans et al., 2012). While all organisms on Earth are constructed from a common set of chemical elements, the relative proportions of carbon (C), nitrogen (N), and phosphorus (P) are important traits that show considerable diversity across ecological contexts and evolutionary lineages (Sterner and Elser, 2002; Kay et al., 2005; Gonzalez et al. 2025). Disentangling the complex hierarchy of factors that drive these fundamental biochemical investments remains a major challenge, as evolutionary history, current ecological opportunities, and intrinsic constraints from physiological processes are all expected to shape organismal composition (Gonzalez et al., 2018; Nessel et al., 2024).

Physiological explanations for stoichiometric variation among organisms include the Growth Rate Hypothesis (GRH), which posits growth-dependent phosphorus allocation to ribosomal RNA (rRNA) in small organisms with high rates of protein synthesis (Elser et al., 1996; Elser et al., 2000) leads to lower C:P and N:P ratios (Isanta-Navarro et al., 2022; Trakimas et al., 2019). Ecological factors are also expected to shape stoichiometry. Aquatic insects, for example, benefit from nutrient-rich organic matter of phytoplanktonic origin, while their terrestrial counterparts must subsist on often nutrient-poor resources and build C-rich structural compounds to counteract gravity (Elser et al., 2000; Shurin, Gruner, and Hillebrand, 2006; Hessen et al., 2013). In addition to differences between aquatic and terrestrial lineages, temperature governs metabolic rates and developmental schedules (Atkinson, 1994; Cross et al., 2015), while nutrient enrichment across successive trophic groups generates stoichiometric mismatches between consumers and their food (Denno and Fagan, 2003; Fagan and Denno, 2004; Wilder and Eubanks, 2010).

Despite an ever-growing body of work on the ecological drivers of stoichiometric variation among taxa (Gonzalez et al. 2025), the influence of evolutionary history on elemental composition remains largely unexplored (Leal et al., 2017). This dearth is significant: association with aquatic or terrestrial ecosystems, metabolic rate, and trophic group are shaped by evolutionary history and covary with phylogeny. Evolutionary history thus may obscure or inflate ecological and physiological drivers, though this has rarely been systematically and robustly tested. Previous studies provide anecdotal evidence for phylogenetic conservatism of stoichiometric traits at relatively small spatial and taxonomic scales (Fagan et al., 2002; Woods et al. 2004, Hendrixson et al., 2006; González et al., 2018), but no study has systematically quantified phylogenetic signal in stoichiometric traits across large, species-rich clades, leaving their macroevolutionary patterns unresolved. Quantifying the relative contributions of ecological, physiological, and evolutionary drivers of stoichiometric variation among species requires an explicitly phylogenetic approach with dense sampling.

Here, we leveraged a global dataset of insect C:N:P stoichiometry spanning 223 species from 13 orders (Gonzalez et al., 2025) and a robust phylogeny (Misof et al., 2014, Tong et al., 2015) to partition the influences of ecology, physiology, and evolution on stoichiometry throughout the insect tree of life (**Fig. 1 a,b)**. After recovering high levels of phylogenetic signal in C:N, N:P, and C:N stoichiometric ratios across insects, we modeled the effects of habitat type (terrestrial, aquatic), mean annual temperature, and body mass on insect stoichiometry using standard (OLS) and evolutionary (PGLS) frameworks. Our approach enables formal comparison between long standing ecophysiological explanations and models that explicitly account for shared ancestry. To quantify the relative influence of evolutionary history on stoichiometric patterns, we partitioned the explanatory power of best-fit models between phylogenetic and non-phylogenetic effects. Finally, we investigated the relative importance of ecology and physiology on body composition by performing phylogenetic PCA (pPCA), which accounts for phylogenetic distance between taxa, followed by regression on pPC scores.

**Figure 1.**
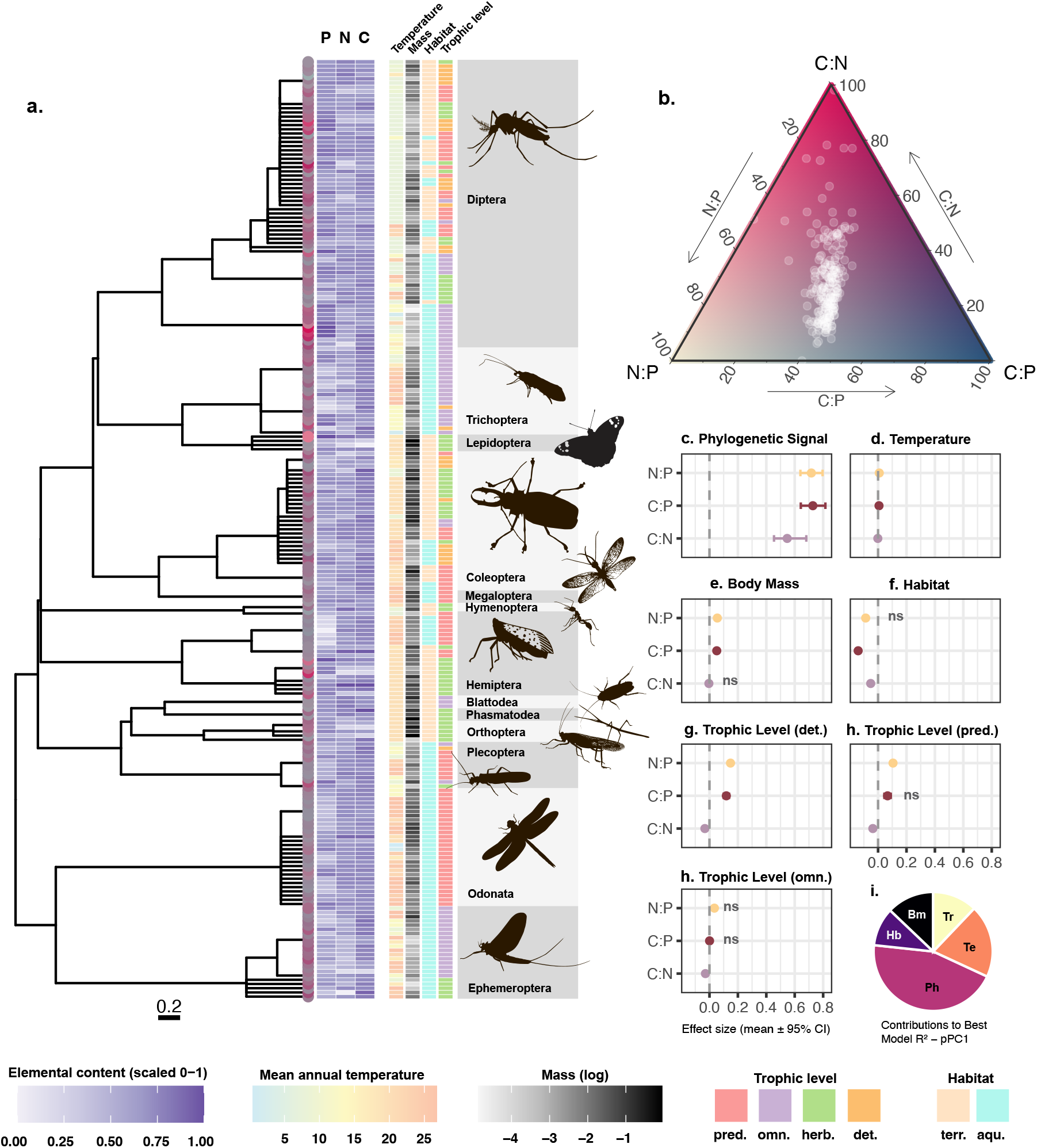
Eco-evolutionary controls of insect stoichiometry. **(a)** The values of C, N, and P, along with important ecological and environmental factors for every species in this analysis plotted against the phylogeny (based on Tong et al. (2015)). **(b)** Ternary plot depicting the scatter in normalized stoichiometric ratios (C:N, N:P and C:P) for all species. **(c)** Lambda-based phylogenetic signal for all stoichiometric ratios. **(d)-(i)** PGLS results showing the effect sizes (with 95% C.I.s) of all ecophysiological (*i*.*e*., non phylogenetic) drivers of stoichiometric variation. **(j)** R^2^ decomposition of the PGLS model for the main axis of variation in pPCA.

We found very strong evidence for phylogenetic signal in all three stoichiometric ratios within insects (**Fig. 1 c)**, with especially strong signals for both phosphorus-based ratios, *i*.*e*. N:P (**λ** = 0.7170; *p* < 10^-5^) and C:P (**λ** = 0.7275; *p* < 10^-5^). C:N composition also showed significant, but weaker, phylogenetic signal (**λ** = 0.5470; *p* < 10^-5^). A multi-trait metric estimated simultaneously for all three ratios likewise recovered significant phylogenetic structure (M = 0.66; *p* < 0.01). These results challenge the idea that organismal stoichiometry is driven solely by ecological factors, and evolutionarily labile, a view implicitly adopted in most previous analyses. Past research on insect stoichiometry was largely unable to detect fine-scale phylogenetic signal, likely due to insufficient phylogenetic depth or differences in sensitivity between estimators (but see González et al., 2018; Hendrixson et al., 2006). Here, we show significant phylogenetic structuring in insect stoichiometric profiles, reasserting the need for an evolutionary perspective.

Accordingly, model selection systematically favored PGLS models with simultaneous estimation of **λ** over conventional regression models (OLS), which themselves outperformed fixed-**λ** (BM) models. R^2^-decomposition of best-fit PGLS models revealed that phylogeny accounts for approximately 48.4% and 45.5% of the total explained variance in C:P and N:P ratios among taxa, respectively, and 40.7% for C:N. Taken together, these results position shared evolutionary history as the principal determinant of insect stoichiometry, with a stronger influence on elemental ratios than any other ecological or physiological predictor (**Fig. 1 j)**.

Even after phylogenetic correction, however, best-fit PGLS models revealed physiological and ecological constraints on insect stoichiometry (**Fig. 1 d-i)**. In line with previously reported patterns of nutrient enrichment across trophic levels (Fagan et al., 2002), detritivores (β = -0.0332; *p* < 0.05) and predators (β = -0.0373; *p* < 0.01) exhibited significantly lower C:N ratios compared to a herbivore baseline. N:P ratios were also higher in these trophic categories than in herbivores (predators: β = 0.105; *p* < 0.01; detritivores: β = 0.150; *p* < 0.01), and detritivore C:P ratios exceeded those of herbivores (β = 0.119; *p* < 0.05).

Body mass was a significant predictor of N:P (β = 0.0526; *p* < 0.01) and C:P (β = 0.0487; *p* < 0.01) ratios, in support of the Growth Rate Hypothesis (Sterner and Elser, 2002). The positive scaling of N:P with body mass confirms that smaller insects maintain lower N:P ratios, consistent with the increased allocation to P-rich rRNA required for rapid growth (Woods et al., 2004).

Echoing past work in algae (Yvon-Durocher et al., 2015) and plants (Sardans et al., 2012), temperature had a positive effect on C:P (β = 0.00705; *p* < 0.01) and N:P (β = 0.00890; *p* < 10^-4^). However, we recovered moderate evidence for a weak negative effect of temperature on C:N (β = -0.00187; *p* < 0.05), a relationship that may have previously evaded detection due to insufficient statistical power (Sardans et al., 2012; Yvon-Durocher et al., 2015). These results may reflect known temperature-dependent declines in nitrogen use efficiency and increases in protein turnover rates in insects (Lemoine and Shantz, 2016). Under this hypothesis, N enrichment of insect tissues in warm climates may support homeostasis by offsetting losses to these processes. Meanwhile, the decrease in P content observed at higher temperatures may be linked to elevated ribosomal efficiency, an established phenomenon in algae (Yvon-Durocher et al., 2015) that may extend to insects.

Finally, relative to aquatic insects, terrestrial insects exhibited lower C:P (β = -0.132; *p* < 0.01 ), C:N (β = -0.0536; *p* < 10^-4^) and marginally lower N:P ratios (β = -0.0775; *p* = 0.057 ). This surprising result contrasts with earlier work (e.*g*., Elser et al., 2000) that reported no difference in the stoichiometry of insects across aquatic and terrestrial habitats. The increased N content in terrestrial insects goes against our original hypothesis that aquatic habitats –which exhibit higher rates of N-fixation per unit-area (Fulweiler et al., 2025)– should support more nutrient-rich organisms. This result, which remains unchanged after accounting for stoichiometric variation across trophic levels, suggests that habitat effects have mechanistic underpinnings that remain unresolved. By contrast, the greater P content of terrestrial insects is likely explained by this element’s sedimentary origin and limiting nature in freshwater ecosystems.

Beyond elemental ratios, pPCA separated two latent dimensions of stoichiometric variation that we respectively interpret as primarily metabolic (pPC1; P) and structural (pPC2; C, N), together explaining 94% of the total variance in insect body composition **(Fig. 1, SX)**. Species placement along pPC1 was significantly influenced by all examined physiological and ecological factors, in line with our analyses of stoichiometric ratios. R^2^ decomposition along this principal axis of variation (66% of total variance) suggested temperature as the most important non phylogenetic determinant of insect stoichiometry (R^2^ = 0.0578), followed by body mass (R^2^ = 0.0337) and trophic level (R^2^ = 0.0332), and finally habitat (R^2^ = 0.0241). Scores along the structural axis (pPC2, 28% of total variance) were predicted by only habitat (β = -0.0563; *p* = 0.00106) in our model.

Our results provide the first evidence for pervasive phylogenetic signal in the elemental composition of insects, challenging current frameworks that do not systematically account for evolutionary distance between taxa. Within these phylogenetic constraints, however, ecological and physiological factors still influence stoichiometry . Our analyses build on and are largely in agreement with some aspects of previous work on the determinants of stoichiometric composition: we highlight strong differences among trophic groups and find support for the growth rate hypothesis. However, we demonstrate N and P enrichment of terrestrial insects relative to aquatic counterparts, contrasting with previous work and hypotheses (Elser et al., 2000; Fulweiler et al., 2025). The key result is that, across the insect clade, phylogeny had an overwhelmingly strong influence on stoichiometric ratios, relative to oft-examined ecological factors (temperature, habitat, trophic level, and body mass). Our approach provides a general template for partitioning evolutionary, ecological, and physiological influences on organismal stoichiometry, and highlights new opportunities to link macroevolutionary patterns to underlying mechanisms.

## Methods

### Stoichiometric and phylogenetic data

We sourced insect stoichiometric data from the StoichLife database (Gonzalez et al., 2025). We restricted our compilation to species with complete cases for elemental composition (%C, %N, %P), dry body mass (g), habitat (aquatic, terrestrial), and mean annual temperature (°C). The resulting dataset comprised 223 species belonging to 14 of the 32 insect orders recognized by Misof et al. (2014). We calculated C:N, N:P, and C:P molar ratios, which were log-transformed alongside elemental compositions to serve as six principal stoichiometric traits.

We modeled phylogenetic relationships between samples by inserting them onto a comprehensive insect backbone by Tong et al. (2015) (itself a reanalysis of the Misof et al. (2014) insect chronogram). Taxa not present in the original tree were placed on the nodes representing the most recent common ancestors of represented sister lineages. Branches restricted to the original tree were pruned, producing a complete phylogeny for our samples. This approach resulted in several polytomies, representing ambiguous relationships between species. To account for this uncertainty, we randomly resolved 500 trees using phytools’ *multi2di* command to serve as inputs for phylogenetically-corrected models. Branch lengths were computed using Grafen’s method (Grafen, 1989) (implemented in the *compute*.*brlen* function of the ape package (Paradis and Schliep, 2019).

### Statistical analysis

We conducted all analyses in R (R Core team, 2025). For all statistical analyses, we treat *p*-values as continuous measures of the strength of evidence, rather than categorical indicators of statistical support.

#### Estimating phylogenetic signal

To identify conserved stoichiometric patterns across the insect tree of life, we first calculated Pagel’s λ (lambda), a standard measure of phylogenetic signal (Pagel, 1999) that was chosen for its robustness to uncertainty in intra-clade phylogenetic distances. We obtained parameter values for all log_10_-transformed traits using the *phylosig* function of the phytools package (Revell, 2024) and averaged them across the 500 replicate trees. For each tree, we assessed the significance of λ using a likelihood ratio test comparing the fit of a model with the estimated λ to a model where λ was fixed at 0. In addition, we used the *phylosignal_M* function of the phylosignalDB package to compute M (Yao and Yuan, 2025), a multi-trait measure of phylogenetic signal that jointly incorporates all three stoichiometric ratios. For all repeated analyses, we report mean parameter estimates and combined *p*-values obtained using aggregated Cauchy association tests –a method robust to non-independent tests– implemented in the ACAT R package.

#### Model fitting, averaging, and selection

To test the impacts of ecological factors and allometry on insect C:N:P stoichiometry, we successively fitted two distinct classes of statistical models: (i) ordinary least-squares (OLS) regressions, which model stoichiometric traits as a linear function of tested predictors and treat species as independent, and (ii) phylogenetic generalized least-squares (PGLS) regression models, which account for the non-independence of species by incorporating the phylogenetic variance-covariance matrix into the model’s error structure (Felsenstein, 1985; Symonds and Blomberg, 2014). We implemented PGLS using the *gls* function in the nlme package (Pinheiro et al., 2017), specifying the phylogenetic correlation structure with *corPagel*. This approach has the advantage of co-estimating the value of Pagel’s λ from the data while fitting the regression model, allowing the model to determine the degree of phylogenetic correction required (Revell, 2024). As an additional check, we also fitted PGLS models assuming pure Brownian motion (implemented in the *corBrownian* function), fixing λ = 1.

We performed PGLS regressions on all 500 replicate trees, followed by model averaging. For all models, we computed the mean and 95% confidence interval of fitted parameter estimates, phylogenetic signal, *t*-values and ACAT *p*-values across all successful fits, *i*.*e*., for all trees on which the regression was successfully performed. Failed model fits were dropped from the analysis.

We based model selection on the Akaike Information Criterion (AIC), with models having the lowest AIC score considered the best fit. For each elemental ratio, we tested the fixed effects of log(body mass), trophic level, habitat and mean annual temperature. We computed AIC tables for each replicate for which all three models (OLS, *corPagel* PGLS, *corBrownian* PGLS) could be fitted. The resulting summary contained, for each model class, the mean and standard deviation of AIC, mean ΔAIC, and the proportion of runs in which it provided the best fit.

#### Phylogenetic PCA (pPCA)

To investigate the integrated evolution of elemental composition, we performed a phylogenetic principal component analysis (pPCA) using the *phyl*.*pca* function of the phytools package while accounting for phylogenetic uncertainty. This analysis was conducted on the covariance matrix of single-element compositions (log(%P), log (%N), log(%C)). pPCA accounts for the phylogenetic relationships among species, enabling the identification of major axes of phylogenetically corrected variation in insect stoichiometry.

To investigate the drivers of sample placement along each principal component, we fitted PGLS or linear-mixed models to pPC scores. For each replicate analysis, phylogenetic signal was evaluated using the *phylosig* function of the phytools package, guiding our selection of model class. For pPC1, which contained residual phylogenetic signal for all replicates, we fitted a PGLS model incorporating habitat, mean annual temperature, log(body mass) and trophic level as fixed effects with *corPagel*. For pPC2, which never displayed phylogenetic signal, we fitted linear models with an identical fixed effect structure. We employed linear model reduction by dropping explanatory variables followed by likelihood ratio tests, as implemented in the *drop1* command of the lme4 package.

#### R-squared decomposition

To quantify the relative contributions of phylogeny and fixed predictors to model fit, we used the *R2_lik* function of the rr2 package (Ives, 2019), which estimates a likelihood-based R^2^ by comparing a candidate model to an appropriate reduced model. We first calculated the overall R^2^ of all best-fit PGLS model using a reduced, non-phylogenetic (OLS) model containing only an intercept. To isolate the non-phylogenetic contribution of model predictors, we then computed R^2^ using a reduced model that retained the phylogenetic structure (PGLS) but excluded all predictors. Finally, we calculated the phylogenetic contribution as the difference between the total R^2^ and the predictor-only component. To further partition the individual contributions of specific ecophysiological factors, we modified this approach by comparing the full model to a series of reduced models that retained the phylogenetic structure (PGLS) and all but the focal predictors.

## Acknowledgements

We thank Nathan Lo for kindly providing the insect chronogram used in this study, and Angélica. L. González and colleagues for making the StoichLife database freely available. Silhouettes from PhyloPic: phasmid (credit: Tree of Life App, CC-BY), plecoptera (credit: Graham Montgomery, CC-BY), mantis (credit: Caleb Gordon, CC-BY); all others public domain.

## Data availability

All code and data for this manuscript are available at https://doi.org/10.5281/zenodo.17904396.

## Author contributions

L.L. led the analyses, wrote the first draft of the manuscript, and developed all final scripts. L.L., R.J.K., and A.S. conceived the study, designed the figures, and contributed to initial conceptual development. All authors contributed to the overall direction of the project and to the final manuscript.

## References

Atkinson, D. (1994). Temperature and organism size – a biological law for ectotherms? Advances in Ecological Research, 25, 1–58.

Elser, J. J., Dobberfuhl, D. R., MacKay, N. A., & Schampel, J. H. (1996). Organism size, life history, and N: P stoichiometry. BioScience, 46(9), 674–684.

Elser, J. J., Fagan, W. F., Denno, R. F., Dobberfuhl, D. R., Folarin, A., Huberty, A., … & Sterner, R. W. (2000). Nutritional constraints in terrestrial and freshwater food webs. Nature, 408(6812), 578–580.

Fagan, W. F., Siemann, E., Mitter, C., Denno, R. F., Huberty, A. F., Woods, H. A., & Elser, J. J. (2002). Nitrogen in insects: implications for trophic complexity and species diversification. The American Naturalist, 160(6), 784–802.

Fagan, W. F., & Denno, R. F. (2004). Stoichiometry of actual vs. potential predator–prey interactions: insights into nitrogen limitation for arthropod predators. Ecology Letters, 7(9), 876–883.

Fulweiler, R. W., Rinehart, S., Taylor, J., Kelly, M. C., Berberich, M. E., Ray, N. E., … & Marcarelli, A. M. (2025). Global importance of nitrogen fixation across inland and coastal waters. Science, 388(6752), 1205–1209.

González, A. L., Merder, J., Andraczek, K., Brose, U., Filipiak, M., Harpole, W. S., … & Aymes, J. C. (2025). StoichLife: A global dataset of plant and animal elemental content. Scientific Data, 12(1), 1–13.

Grafen, A. (1989). The phylogenetic regression. Philosophical Transactions of the Royal Society of London. B, Biological Sciences, 326(1233), 119–157.

Hessen, D. O., Elser, J. J., & Sterner, R. W. (2013). Ecological stoichiometry: an elementary approach using basic principles. Limnology and Oceanography, 58(6), 2219–2236.

Isanta-Navarro, J., Prater, C., Peoples, L. M., Loladze, I., Phan, T., Jeyasingh, P. D., … & Elser, J. J. (2022). Revisiting the growth rate hypothesis: Towards a holistic stoichiometric understanding of growth. Ecology Letters, 25(10), 2324–2339.

Ives, A. R. (2019). R s for correlated data: Phylogenetic models, lmms, and glmms. Systematic Biology, 68(2), 234–251.

Leal, M. C., Seehausen, O., & Matthews, B. (2017). The ecology and evolution of stoichiometric phenotypes. Trends in Ecology & Evolution, 32(2), 108–117.

Lemoine, N. P., & Shantz, A. A. (2016). Increased temperature causes protein limitation by reducing nitrogen digestion efficiency in the ectothermic herbivore Spodoptera exigua. Physiological Entomology, 41(2), 143–151.

Misof, B., Liu, S., Meusemann, K., Peters, R. S., Donath, A., Mayer, C., … & Zhou, X. (2014). Phylogenomics resolves the timing and pattern of insect evolution. Science, 346(6210), 763–767.

Pagel, M. (1999). Inferring the historical patterns of biological evolution. Nature, 401(6756), 877–884.

Paradis, E., & Schliep, K. (2019). ape 5.0: an environment for modern phylogenetics and evolutionary analyses in R. Bioinformatics, 35(3), 526–528.

Pinheiro, J., Bates, D., DebRoy, S., Sarkar, D., Heisterkamp, S., Van Willigen, B., & Maintainer, R. (2017). Package ‘nlme’. Linear and nonlinear mixed effects models, version, 3(1), 274.

R Core Team (2025). R: A Language and Environment for Statistical Computing. R Foundation for Statistical Computing, Vienna, Austria. https://www.R-project.org/.

Revell, L. J. (2024). phytools 2.0: an updated R ecosystem for phylogenetic comparative methods (and other things). PeerJ, 12, e16505.

Sterner, R. W., & Elser, J. J. (2002). Ecological stoichiometry: the biology of elements from molecules to the biosphere. Princeton University Press.

Symonds, M. R., & Blomberg, S. P. (2014). A primer on phylogenetic generalised least squares. In Modern phylogenetic comparative methods and their application in evolutionary biology (pp. 105–130). Springer Berlin Heidelberg.

Tong, K. J., Duchêne, S., Ho, S. Y. W., & Lo, N. (2015). Comment on “Phylogenomics resolves the timing and pattern of insect evolution”. Science, 349(6247), 487–487.

Woods, H. A., Fagan, W. F., Elser, J. J., & Harrison, J. F. (2004). Allometric and phylogenetic variation in insect phosphorus content. Functional Ecology, 18(1), 103–109.

Yao, L., & Yuan, Y. (2025). A Unified Method for Detecting Phylogenetic Signals in Continuous, Discrete, and Multiple Trait Combinations. Ecology and Evolution, 15(3), e71106.

Yvon-Durocher, G., Dossena, M., Trimmer, M., Woodward, G., & Allen, A. P. (2015). Temperature and the biogeography of algal stoichiometry. Global Ecology and Biogeography, 24(5), 562–570.

